# Beak elongation as a candidate key innovation underlying ecological transitions in shorebirds

**DOI:** 10.1101/2025.08.24.671984

**Authors:** Michel Baguette, Glenn Le Floch, Eleanor Lacassagne, Vincent Bels, Nicolas Schtickzelle

**Author notes:** Co-first authors (Michel Baguette, Glenn Le Floch).

## Abstract

Adaptive radiations are often associated with phenotypic innovations that broaden ecological opportunity and bias subsequent ecological divergence. In shorebirds (Charadriiformes), variation in beak length has long been linked to resource partitioning, yet its role as a functional innovation has remained largely untested. Here, we integrate behavioural, morphological, and phylogenetic data from 29 Western Palearctic shorebird species to evaluate whether beak elongation fulfils key expectations of a candidate innovation by being repeatedly associated with ecological and functional transitions. Species with longer than expected beaks relative to body size consistently exhibit distinct locomotor, prey-capture, and food-transport strategies. These integrated morphological-behavioural syndromes recur across independent lineages, suggesting convergent evolution driven by shared ecological constraints. Phylogenetic analyses indicate multiple independent origins of relative beak elongation, supporting its role as a repeated evolutionary solution. Together, our results support a functionally integrated view of adaptation, in which coordinated changes in morphology, sensory investment, and behaviour facilitate access to alternative foraging microhabitats and enable repeated ecological transitions within shorebirds. More broadly, this study provides a testable framework for identifying candidate innovations based on replicated trait-niche associations.

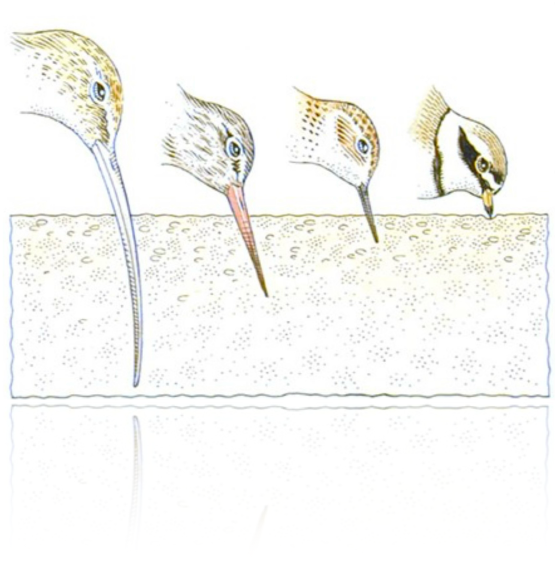

## Introduction

Adaptive radiation refers to the evolutionary emergence of phenotypic diversity within a lineage as a response to divergent selection pressures (Schluter 2000; Gavrilets & Losos 2009). This process often depends on the availability of novel ecological niches, which may be accessed through the emergence of a key innovation - that is, a phenotypic modification enabling individuals to exploit previously inaccessible ecological opportunities (Stroud & Losos 2016). While the role of key innovations in driving adaptive radiation has been widely acknowledged, Miller et al. (Miller et al. 2023) argue that the concept has gradually shifted in meaning. Specifically, they note a conflation between lineage diversification (e.g., Simões et al. 2016) and trait-dependent ecological transitions (e.g., Baguette et al. 2020). In response, Miller et al. (Miller et al. 2023) advocate for a return to a more ecologically grounded definition of key innovations. They propose two complementary approaches to operationalizing this concept, by elucidating the mechanisms through which phenotypic traits alter the interaction between organisms and their environments, and/or demonstrating that these trait changes lead to measurable shifts in the organism’s ecological niche. In this study, we adopt the second of the two approaches proposed by Miller et al. (Miller et al. 2023), focusing on identifying phenotypic shifts that may have facilitated facilitated ecological transitions in shorebirds.

Shorebirds represent a relatively phenotypically homogeneous group of approximately 215 species (Colwell 2010), encompassing two of the three suborders within the order Charadriiformes: Scolopaci (e.g., shanks, tattlers, sandpipers, godwits, snipes, curlews) and Charadrii (e.g., plovers, avocets, oystercatchers, lapwings). A recent time-calibrated phylogeny - integrating genetic, morphological, and paleontological data - provides a robust framework for understanding the remarkable species diversity found within the Charadriiformes (Černý & Natale 2022). Shorebirds constitute one of the most prominent avian radiations, not only in terms of species richness but also with respect to morphological, behavioral, and biogeographical variation (van Tuinen et al. 2004; Colwell 2010,). They are frequently cited as a textbook example of resource partitioning, wherein contrasting morphological traits - particularly beak and leg structures - facilitate the localization, capture, and ingestion of food items. During the non-breeding season, most shorebirds predominantly forage along land - water interfaces in freshwater, brackish, or marine environments, where they feed primarily on invertebrates, but also on small vertebrates and plant material. It is widely assumed that interspecific variation in beak length allows different shorebird species to access prey located at different sediment or water column depths, while variation in leg length permits foraging across a range of water depths. However, the functional and ecological correlates of beak-length divergence have rarely been evaluated quantitatively in an explicit comparative framework.

In a previous study, we described and classified the food acquisition behaviors of 26 western Palearctic shorebird species during migration and wintering periods (Figure 1 in [Baguette et al. 2024]). Among these, 8 species belonged to the suborder Charadrii and 18 to Scolopaci. This work revealed the existence of a behavioral syndrome associated with food acquisition, characterized by covariation among stereotyped behaviors used during each of the three fundamental steps of foraging: food detection, capture, and transport from the tip of the beak to the pharynx. Furthermore, our findings suggested a phylogenetic component to this syndrome, species with similar behavioral covariation cluster together in the phylogeny. We also demonstrated that each of these three behavioral steps constitutes an ecological performance in the sense of Irschick & Higham (Irschick & Higham 2016), *i*.*e*., quantifiable traits that reflect how effectively an individual performs tasks critical to its fitness.

**Fig. 1.**
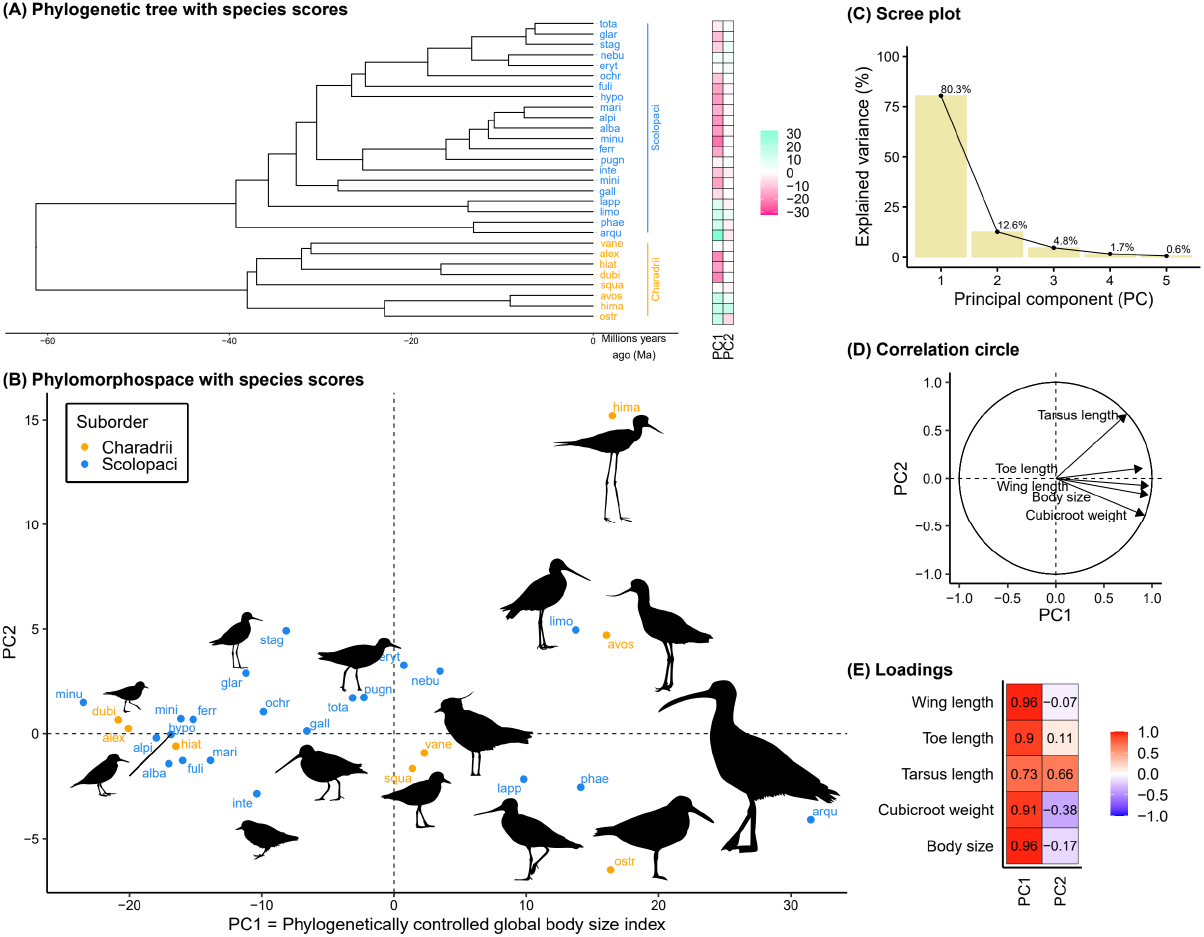
Shorebird phylomorphospace. We used the phylogenetic tree of the 29 shorebirds species (**A**) to perform a phylogenetic constrained Principal Component Analysis (pPCA) on five morphological traits and create a phylomorphospace (**B**) using its first two principal components (PC1 and PC2). PC1 summarizes 80% of the variance (**C**) and can be interpreted as a phylogenetically constrained global body size index, being highly and positively correlated with the five traits (**D**) and (**E**). 14 species are represented on the phylomorphospace (**B**) at the same scale (bird silhouettes from phylopic.org; detailed credits in the supplementary material: code S1 and data S3).

In the present study, we expand our behavioral dataset by incorporating information from three additional Scolopaci species. In parallel, we compiled morphological data related to body structure (body mass, body length, and the size of musculoskeletal appendages), sensory organs (eye diameter and the number of sensory pits on the beak - serving as proxies for visual and tactile foraging abilities, respectively), as well as beak and tongue lengths. To develop a standardized morphological baseline, we first combined the body structure traits (excluding beak length) to construct a global body size index for each of the 29 species, using a phylogenetically informed approach. We then quantified allometric relationships between this index and key morphological traits involved in food acquisition - specifically, sensory structures, beak, and tongue - representing their roles in food detection, capture, and transport, respectively. Using these relationships, we calculated Relative-to-Expected (RTE) trait values, defined as species-specific deviations of a given morphological trait from its predicted value based on the global body size index. To test potential differences in evolutionary trajectories between the two shorebird suborders, we compared RTE trait values between Charadrii and Scolopaci. Lastly, we investigated whether, and in what manner, the three components of the food acquisition behavioral syndrome - locomotion, prey capture, and transport of the food from the tip of the beak to the pharynx - are associated with deviations in RTE beak length across species.

## Material and Methods

### Phenotypic traits

We collected data on the food acquisition behavior of 29 species in wintering or migration sites (Baguette et al. 2024; Hernandez--Possémé et al. in review), which represents ca. 75% of the total shorebird fauna of the western Palearctic (Cramp & Simmons 1983, except for vagrants), and ca. 90% of the species using land - water interfaces during migration and overwintering (Cramp & Simmons 1983, except for vagrants). Locomotion behaviors (how individuals move around their feeding habitats), capture behaviors (how individuals catch and locate their prey) and transport behaviors (how individuals move food particles from the tip of the beak to the pharynx) were categorized based on a recent repertoire (Baguette et al. 2024). There are three capture behaviors (pecking, probing, pecking-probing), two locomotion behaviors (stop-run-stop, continuous walking), and three transport behaviors (lingual, ballistic and ballistic-surface tension).

Aggregated data at the species level were used for morphological variables (beak length, body size, weight, wing length, tarsus length, length of the longest toe, tongue length) and sensory organs (diameter of the eye, number of sensory pits on the beak). Because behavioural categories were assessed at the species level, our analyses explicitly target macroevolutionary associations among species. They are therefore not designed to capture within-species behavioural reaction norms or individual-level plasticity, but rather to identify replicated patterns of trait - behaviour covariation across lineages. Morphological data come from Cramp & Simmons (Cramp & Simmons 1983), except tongue lengths, which were published by Burton (Burton 1974), diameters of the eye that come from (Thomas et al. 2006) and the number of sensory pits on the beak that were measured by one of us (EL) on the available specimens of the Compared Anatomy Lab of the Muséum National d’Histoire Naturelle (Paris). The values used are averages across sexes, and when values from different geographical origins were available, we used those measured in the localities closest to our behavior observation sites. Four variables were available for a subset of species only: eye diameter (22 species), number of sensory pits (9 species) and tongue length (19 species). Two variables were transformed before analysis: weight was cubic root transformed to account for its scaling as a volume, not a length; tongue length was expressed as a proportion of the beak length and this tongue/beak length ratio was ln transformed to make its distribution symmetric.

### Statistical analyses

First, we summarized the variation among the 29 species in five morphological traits (body size, wing length, tarsus length, length of the longest toe, and cubic root of weight) using a phylogeny constrained Principal Component Analysis (PCA; phyl.pca function from *phytools* R package) (14,15) to control for the effect of evolutionary relatedness among species. Phylogenetic distances used in this PCA were inferred using a pruned tree restricted to our study species from the Charadriiformes time tree of Černý & Natale (Černý & Natale 2022). The first two Principal Components (PCs) were used to place the species in a phylomorphospace. PC1 summarizes a very high proportion (80%) of the morphological variation and can be interpreted as a phylogenetically controlled global body size index because it is highly and positively correlated with all the five morphological traits.

Second, we used this phylogenetically controlled global body size index to correct morphological traits of interest (eye diameter, number of sensory pits, beak length, and tongue/beak length ln-ratio) for allometry. A linear regression between the trait and the body index was estimated (lm function from *stats R* package) (R Core Team 2025), and a new Relative-To-Expected (RTE) trait variable was created to express how much a species possesses a trait value that differs from its expected value given its body size index; RTE trait for a species was computed as the *ln* transformed ratio between its residual value and its predicted value, both from the allometric regression. We tested whether species from the two suborders significantly differ for these four RTE traits using two sample t-tests.

Third, we checked for specific relationships among morphological traits and behaviors associated to the three main steps of food acquisition (locomotion, capture and transport) using appropriate linear models according to the nature of the response variable: linear regression (lm function from *stats R* package) (R Core Team 2025) for a continuous normal variable, binomial logistic regression (glm function from *stats R* package) (R Core Team 2025) for a two levels discrete variable, and binomial logistic regression (vglm function from *VGAM R* package) for an ordered multilevel discrete variable.

## Results

We constructed a phylomorphospace (Figure 1) by plotting species scores along the first two axes of a phylogenetically constrained Principal Component Analysis (pPCA) (Revell 2009; Revell 2012), which summarizes interspecific variation in five morphological traits across the 29 studied shorebird species. The first principal component (PC1), which accounts for 80% of the total morphological variation, was interpreted as a phylogenetically controlled global body size index. Along this axis, species range from the smallest plovers and sandpipers to the largest taxa, such as the avocet, stint, godwit, and curlew, with the largest species being 3.8 times larger than the smallest.

Among the traits analyzed, eye diameter increases significantly with body size. After correcting for this allometric relationship, RTE eye diameter is significantly larger in Charadrii than in Scolopaci (Figure 2A). In contrast, the number of sensory pits on the beak decreases significantly with body size and, when corrected for allometry, is significantly higher in Scolopaci than in Charadrii (Figure 2B). A negative correlation between the two RTE sensory traits indicates that species with fewer sensory pits tend to have relatively larger eyes (Figure 3). Beak length scales positively with body size, and once corrected for allometry, species in Charadrii have significantly shorter beaks than those in Scolopaci (Figure 2C). Additionally, the ratio of tongue length to beak length shows a significant negative scaling relationship with body size: larger species possess proportionally shorter tongues, although not necessarily in absolute length. However, after correcting for this allometric effect, no significant difference in tongue-to-beak length ratio is observed between the two suborders (Figure 2D).

**Fig. 2.**
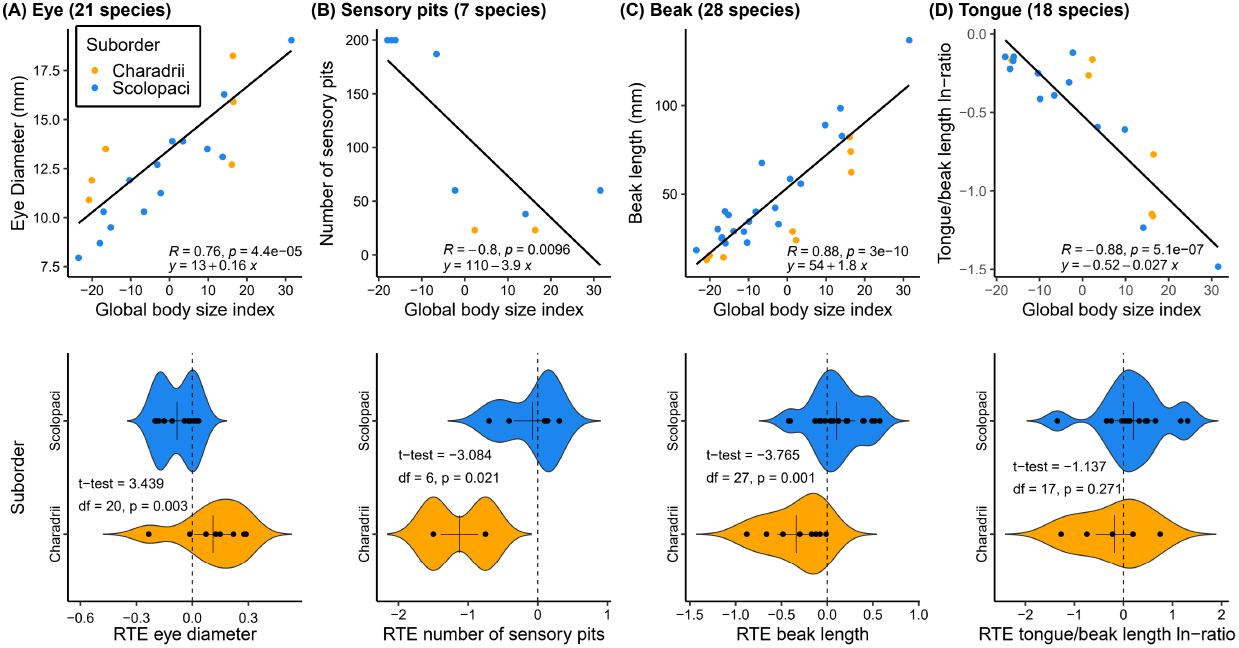
Allometric relationships. We use allometric relations (first line, with linear regression statistics) with the phylogenetically controlled global body size index to correct four morphological traits for allometry (i.e. the species body size index) by computing their RTE (Relative-to-Expected) counterparts. We then check whether these RTE traits differ between the two suborders, Charadrii and Scolopaci (second line, with t-test statistics). The four morphological traits are the eye diameter (A), the number of sensory pits on the beak (B), the beak length (C) and the tongue length in proportion of the beak, expressed as the ln-ratio of tongue over beak length (D). Sample size for each trait is indicated in panel titles.

**Fig. 3.**
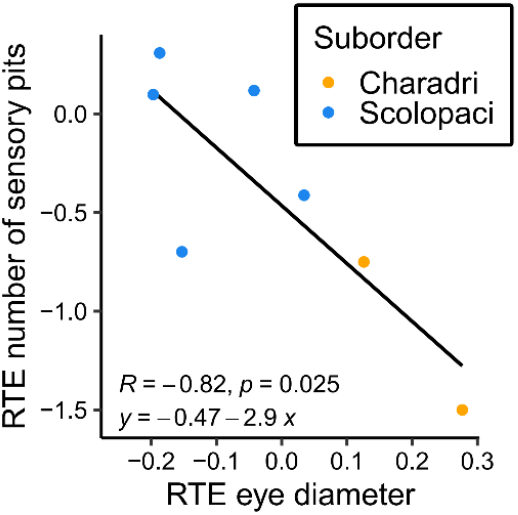
Relationship between eye diameter and sensory pits. Species with a smaller eyes have more sensory pits on their beak. Both variables are corrected for their allometric relation with body size, being expressed as Relative-to-Expected (RTE) values. Statistics refer to a linear regression.

Relative-to-Expected (RTE) beak length is a significant predictor of behavioral strategies employed during food acquisition. Species with longer than expected beaks (relative to their body size index) are more likely to exhibit: (1) continuous walking rather than stop-run-stop locomotion (Figure 4A); (2) probing rather than pecking as a prey capture technique, with transitional species exhibiting a combination of both behaviors (Figure 4B); and (3) ballistic rather than lingual mechanisms for food transport, with transitional species employing both surface tension and ballistic strategies (Figure 4C). In contrast, the relative tongue length (corrected for beak length) does not significantly influence the mode of food transport behavior (Figure 4D).

**Fig. 4.**
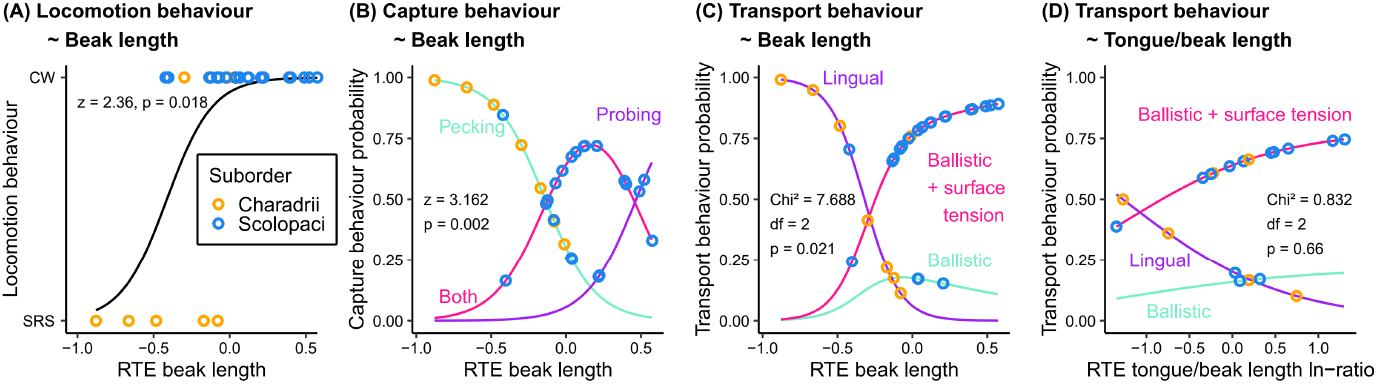
Relationship between beak/tongue length and food acquisition behaviors. Locomotion (A), prey capture (B) and food transport (C) behaviors used by a species significantly depend on how longer than expected its beak is given its body size index. Transport behavior is however not influenced by how longer its tongue (relative to beak length) is (D). Beak length and the ration tongue/beak length are corrected for their allometric relation with body size, being expressed as Relative-to-Expected (RTE) values. Continuously walking (CW) and stop-run-stop (SRS) are the main two locomotor modes used by shorebirds (*12,20*). Statistics refer to binomial (A), cumulative (B) or multinomial (C and D) logistic regressions.

## Discussion

Outside the breeding season, shorebird species tend to aggregate in large numbers within suitable foraging habitats, predominantly along coastal and shoreline environments (*e*.*g*., van der Kam et al. 2004; Colwell 2010). On overwintering grounds, food biomass is limited (Lourenço et al. 2018); moreover, for prey items to be considered profitable, they must be accessible, detectable, manageable (i.e., handleable and ingestible), and digestible (van der Kam et al. 2004). Consequently, interspecific food partitioning within shorebird communities is commonly observed (*e*.*g*., Catry et al. 2016). Recent advances in stable isotope analysis and DNA metabarcoding have provided important insights into the dietary composition of co-occurring shorebird species (Catry et al. 2016; Correia et al. 2023). These studies reveal that species often exhibit local and temporal dietary specializations, frequently relying on specific prey types, and developing cues and behaviors tailored to exploiting those prey efficiently. A substantial body of research grounded in optimal foraging theory has demonstrated that foraging shorebirds tend to maximize net energy gain by optimizing prey selection, handling, and ingestion strategies (*e*.*g*., van der Kam et al. 2004; Colwell 2010). However, despite these insights into behavioral decision-making, questions remain regarding the evolutionary processes that have shaped the morphological and behavioral traits underlying such ecological specialization.

Sensory organs play a critical role in the successful detection of food. Shorebirds employ two primary strategies for locating prey or plant material: visual detection (hunting by sight) and tactile detection (hunting by touch), with some species potentially combining both approaches (*e*.*g*., van der Kam et al. 2004; Baguette et al. 2024; Barbosa & Moreno 1999; Thomas et al. 2006; Lourenço et al. 2016; du Toit 2021; Martin 2017). In the present study, we use eye diameter and the number of sensory pits on the beak as proxies for a species’ relative investment in visual and tactile prey detection, respectively. Across the 29 species in our dataset, eye diameter scales positively with the global body size index. Notably, six of the eight Charadrii species exhibit eye diameters that are - sometimes substantially - larger than expected based on this allometric relationship. According to the dated phylogeny by Černý and Natale (Černý & Natale 2022), our Charadrii sample includes five species from the Charadriidae family (plovers and lapwing), which represents an early-diverging clade within the suborder, and three species from more recently diverged lineages. The six species with larger-than-expected eye diameters comprise the five Charadriidae representatives and the oystercatcher, suggesting that enlarged eye size is a retained trait within the early-diverging clade.

In contrast, the number of sensory pits on the beak shows a significant negative relationship with the global body size index. However, sensory pit data are lacking for many species, particularly within Charadrii, for which data are available for only two species: the lapwing (representing the basal clade) and the oystercatcher (a representative of the more recently diverged clade). Conclusions regarding sensory investment should therefore be interpreted cautiously given limited taxonomic coverage for sensory pits, and we treat them as preliminary proxies rather than definitive measures. Despite these data limitations, the observed differences in sensory organ morphology between Charadrii and Scolopaci are broadly consistent with their known prey capture strategies. Most Charadrii, especially those from the basal clade, exhibit larger than expected eye diameters and rely predominantly on visual prey detection. In contrast, Scolopaci species, which locate prey by touch or through a combination of tactile and visual cues, possess more sensory pits on their beaks than expected given their body size. Furthermore, although based on a limited sample, our findings suggest a potential trade-off in sensory investment: species with larger-than-expected eye diameters tend to exhibit a lower than expected number of sensory pits, implying a specialization towards either visual or tactile foraging, but rarely both.

In our dataset, beak length exhibits a positive allometric relationship with the global body size index, indicating that larger shorebird species generally possess longer beaks. However, after correcting for this allometric effect, species belonging to the suborder Charadrii display significantly shorter than expected beak lengths compared to those in Scolopaci. This pattern is primarily driven by the five species within the family Charadriidae (plovers and lapwing), the early-diverging clade within Charadrii. In contrast, species from the more recently diverged (Haematopodidae + Recurvirostridae) clade (e.g., oystercatchers, avocets, and stints) exhibit relative beak lengths comparable to those of Scolopaci species.

The tongue plays a crucial role in the evolution of food transport mechanisms in Sauropsida, including birds (Bels et al. 2023). Among shorebirds, tongue length varies markedly, ranging from nearly equal to beak length to less than one-third of it (Burton 1974). To account for this diversity, we expressed tongue length as a proportion of beak length and found that this ratio scales negatively with body size. These findings suggest that tongue elongation is subject to an upper constraint that prevents proportional increases in length relative to the beak, possibly due to biomechanical or physiological trade-offs. Such constraints may be linked to the evolution of specific food transport mechanisms, such as ballistic transport (Baussart et al. 2009; Harte et al. 201; Bels et al. 2023) or surface tension-based transport (Rubega & Obst 1993; Rubega 1997; Prakash et al. 2008; Bels et al. 2023,29-31).

In previous work, we decomposed shorebird food acquisition behavior into three sequential stages: foraging, feeding, and swallowing (12). Foraging encompasses locomotion and prey capture - respectively focused on locating and seizing food items - while feeding refers to the manipulation and transport of food toward the beak. Swallowing marks the final stage of food acquisition, when food is ingested into the pharynx. Locomotion, capture, and transport are three performances (13) that can be further subdivided into three to four stereotyped behaviors, following Tinbergen’s hierarchical model of behavioral complexity (32). These behaviors are readily observable and quantifiable under field conditions (12).

We previously identified a behavioral syndrome in shorebirds characterized by consistent associations among stereotyped behaviors of each of these performances (Baguette et al. 2024). Pecking is considered the plesiomorphic prey capture behavior in shorebirds (Zweers 1991; Zweers & Gerritsen 1997), and all five Charadriidae species in our dataset exhibit a behavioral profile characterized by stop-run-stop locomotion, pecking as the primary capture strategy, and lingual transport of food particles (Baguette et al. 2024). In contrast, species from the Scolopaci suborder and from the more recently diverged Charadrii clade (Haematopodidae + Recurvirostridae) employ a mixed capture strategy combining pecking and probing, use continuous walking as their primary locomotion behavior, and - except for one species - utilize a combination of ballistic and surface tension-based food transport strategies.

Here, we demonstrate that this behavioral syndrome aligns closely with differences in relative beak length between the basal and more derived clades. Species with shorter than expected beaks relative to body size consistently employ the behavioral combination of stop-run-stop locomotion, pecking capture, and lingual transport. Conversely, species with longer than expected beaks are more likely to exhibit probing or combined pecking-probing behaviors, continuous walking, and employ ballistic or surface tension transport mechanisms. These findings support the hypothesis that morphological divergence in beak length has co-evolved with functional behavioral adaptations that optimize food acquisition strategies within distinct ecological contexts.

In particular, the convergence of morphological and behavioral phenotypes observed between long-beaked Scolopaci and the recently diverged long-beaked Charadrii further supports the idea that beak elongation is repeatedly associated with shifts in locomotion, prey capture, and food transport, consistent with a functional role in enabling alternative food-acquisition strategies. The beak, serving as the primary grasping organ in birds and often analogized to the primate hand (Bhullar et al. 2016), plays a fundamental role in food acquisition (Xu et al. 2023). However, studies examining the relationship between beak morphology and feeding ecology across modern birds as a whole have yielded inconclusive results (Navalón et al. 2019). Two principal factors likely account for this lack of a clear relationship: first, similar trophic resources may be exploited via markedly different beak morphologies; second, beak morphology may be shaped by selective pressures unrelated to feeding, such as communication, nest construction, preening, or thermoregulation (Olsen 2017). These confounding factors can be mitigated by focusing analyses at lower taxonomic levels (Navalón et al. 2019), as we do here, and as previously demonstrated by Olsen (Olsen 2017), who found feeding ecology to be the primary driver of beak diversification within Anseriformes.

Our dataset encompasses most shorebird species that forage along land - water interfaces during migration and overwintering in the Western Palearctic. During this period, individuals concentrate in spatially restricted habitats where prey accessibility and detectability can be limiting, potentially amplifying functional differences among co-occurring species. Importantly, much diversification within several shorebird genera has been linked to late Cenozoic climatic dynamics (Černý & Natale 2022), whereas the evolutionary emergence of pronounced beak elongation likely predates many of these more recent lineage splits. In this context, we propose that beak elongation represents a strong candidate innovation at the functional - ecological level: it is repeatedly associated with coordinated shifts in sensory investment and in the behavioural repertoire underlying food acquisition. This pattern is consistent with the view that certain phenotypic transitions can bias subsequent ecological divergence by enabling lineages to exploit alternative foraging microhabitats or prey-access pathways, even if they do not directly translate into detectable diversification-rate shifts. Our study therefore supports an explicitly testable framework based on replicated trait-niche associations for evaluating candidate innovations in shorebirds and beyond.

One might question whether this hypothesis is merely a “just-so” story, as criticized by Gould and Lewontin (Gould & Lewontin 1979) for many adaptive explanations. As with most historical evolutionary hypotheses, our inference relies on comparative evidence rather than direct experimental reconstruction. Nonetheless, the repeated association between relative beak elongation and a tightly integrated food-acquisition syndrome across independent lineages is difficult to reconcile with coincidence alone and is consistent with convergent selection under similar ecological constraints. Future work combining broader taxon sampling and direct ecological measures (e.g., water depth use, prey accessibility, and fine-scale diet composition when informative) will help refine the mechanistic links between beak elongation and niche transitions.

Altogether, our findings provide empirical support for the idea that morphological innovations such as beak elongation can drive the emergence of novel behavioral repertoires, thereby opening new ecological opportunities. This functional integration between structure and behavior highlights a broader principle in evolutionary ecology: innovations are rarely isolated events, but rather part of coordinated phenotypic transitions. While the concept of “key innovations” is often invoked post hoc, our results point to a specific sequence - elongation of the beak, followed by shifts in sensory investment and feeding mechanics - as plausible evolutionary pathway underpinning niche differentiation in shorebirds. This underscores the need to consider behavioral and sensory traits in tandem with morphology when evaluating the drivers of adaptive radiation. Furthermore, the convergence observed between unrelated long-beaked lineages suggests strong selective constraints and offers a rare example of replicated evolutionary solutions within a single avian order.

## Supporting information

Sup_Mat

## Acknowledgments

We warmly thank Hans Van Dyck and Joris Bertrand for constructive comments, Raphaël Cornette and Frédéric Legendre for helpful methodological advice, and Virginie M. Stevens, Delphine Legrand and Hervé Philippe for discussions and daily support. Lucyle Hernandez -- Possémé granted access to unpublished results. This manuscript is publication BRCXXX of the Biodiversity Research Centre at UCLouvain, to which N.S. is affiliated.

## Funding

Internship UMR 7205 (EL)

Doctoral fellowship ED226, MNHN (GLF) Internal grants UMR 7205 (MB, VB)

Agence Nationale de la Recherche grant TULIP 10-LABX-0041 (MB)

## Author contributions

Conceptualization: MB

Study design MB, GLF, VB, NS

Data Acquisition: MB, GLF, EL

Methodology: MB, NS, VB

Data Analysis: NS Visualization: NS

Supervision: MB

Writing - original draft: MB

Writing - review & editing: MB, GLF, EL, VB, NS

## Competing interests

Authors declare that they have no competing interests.

## Data and materials availability

All data and code are available in the supplementary material.

## Notes

### Competing Interest Statement

The authors have declared no competing interest.

### Summary of Updates

In this revised version of the manuscript, we have implemented a set of targeted refinements aimed at clarifying the scope of the study, strengthening the alignment between data and interpretation, and improving overall coherence. These changes were motivated by constructive scientific discussions with colleagues and by our own efforts to sharpen the ecological and comparative focus of the work. First, we have clarified the conceptual framing of the study by explicitly treating beak elongation as a candidate key innovation in an ecological and functional sense. In line with recent theoretical work, we now emphasize replicated trait-niche associations and repeated ecological transitions, rather than making claims about diversification rates per se. This adjustment better reflects both the structure of our dataset and the questions addressed by our analyses. Second, we have streamlined the evolutionary context in which our results are discussed. Interpretations extending beyond the temporal and taxonomic scope of the data have been reduced or reformulated in a more cautious manner. In particular, broader deep-time scenarios are no longer used as central explanatory elements, ensuring that evolutionary inferences remain closely anchored to the comparative evidence presented here. Third, the manuscript has been reorganized to improve clarity and transparency. The Introduction now more clearly separates background, rationale, and objectives. The Methods section explicitly defines the level of biological organization addressed by the analyses. In particular, we now state that behavioural categories were assessed at the species level, and that the analyses therefore target macroevolutionary associations among species rather than within-species behavioural reaction norms or individual-level plasticity. This clarification helps to delimit the inferential scope of the study and avoid potential misinterpretations. Fourth, the Results and Discussion have been edited to strengthen the correspondence between empirical patterns and their interpretation. Greater emphasis is placed on the repeated association between relative beak elongation and coordinated shifts in locomotion, prey capture, and food transport across independent lineages. This highlights convergence and functional integration, rather than singular evolutionary events, and reinforces the generality of the conclusions. Overall, these revisions were designed to improve precision, robustness, and readability while preserving the core empirical findings and their ecological significance. We believe that the revised manuscript more accurately reflects the strengths and intended scope of the study, and that it provides a clearer and more testable contribution to discussions on functional innovations and ecological transitions in adaptive radiations.

