## Supplementary material for "Beak elongation as a candidate key innovation underlying ecological transitions in shorebirds": Sup_Mat

Michel Baguette *et al.*

**This PDF file includes:**

Code S1
Data S1, S2
Reference (1)

**Other Supplementary Materials for this manuscript include the following:**

Data S1, S2

**Code S1.**

R script producing all results and figures presented in the manuscript. Requires data files S1 to S3.

```
##########################################################################
## Production of results and figures of paper                           ##
## Baguette, Le Floch, Lacassagne, Bels & Schtickzelle                   ##
## Is beak elongation a key innovation for shorebird adaptive radiation?  ##
## Written by Nicolas Schtickzelle (UCLouvain)                          ##
##########################################################################


## To run this script, please specify the working directory where you placed
##  * the datasets (shorebird_data.csv, shorebirds_tree.tre)
##  * the "Species_silhouettes" subdirectory

setwd ("YOUR_WORKING_DIRECTORY")



# 0. Packages -----------------------------------------------------------
# Install the packages if they are not already installed, then load them
if ( require("data.table")   == FALSE ) install.packages("data.table")
# Extension of `data.frame`
 library("data.table")
if ( require("dplyr")        == FALSE ) install.packages("dplyr")
# A Grammar of Data Manipulation
 library("dplyr")
if ( require("RColorBrewer") == FALSE ) install.packages("RColorBrewer")
# ColorBrewer Palettes
 library("RColorBrewer")
if ( require("ggplot2")      == FALSE ) install.packages("ggplot2")
# Create Elegant Data Visualisations Using the Grammar of Graphics
 library("ggplot2")
if ( require("ggforce")      == FALSE ) install.packages("ggforce")
# Accelerating 'ggplot2'
 library("ggforce")
if ( require("ggrepel")      == FALSE ) install.packages("ggrepel")
# Automatically Position Non-Overlapping Text Labels with 'ggplot2'
 library("ggrepel")
if ( require("ggpubr")       == FALSE ) install.packages("ggpubr")
# 'ggplot2' Based Publication Ready Plots
 library("ggpubr")
if ( require("remotes")      == FALSE ) install.packages("remotes")
# R Package Installation from Remote Repositories, Including 'GitHub'
 library("remotes")
if ( require("svgparser")    == FALSE ) remotes::install_github('coolbutuseless/
svgparser') # Read SVG as Grobs and Data Frames
 library("svgparser")
if ( require("reshape2")     == FALSE ) install.packages("reshape2")
# Flexibly Reshape Data: A Reboot of the Reshape Package (Flexibly restructure
and aggregate data using just two functions: melt and cast.)
 library("reshape2")
if ( require("rstatix")      == FALSE ) install.packages("rstatix")
# Pipe-Friendly Framework for Basic Statistical Tests
 library("rstatix")
if ( require("FactoMineR")   == FALSE ) install.packages("FactoMineR")
# Multivariate Exploratory Data Analysis and Data Mining
 library("FactoMineR")
if ( require("factoextra")   == FALSE ) install.packages("factoextra")
# Extract and Visualize the Results of Multivariate Data Analyses
 library("factoextra")
```

```
if ( require("ape")          == FALSE ) install.packages("ape")
# Analyses of Phylogenetics and Evolution
 library("ape")
if ( require("phytools")     == FALSE ) install.packages("phytools")
# Phylogenetic Tools for Comparative Biology (and Other Things)
 library("phytools")
if ( require("BiocManager")  == FALSE ) install.packages("BiocManager")
# Access the Bioconductor Project Package Repository
 if ( require("ggtree")      == FALSE ) BiocManager::install("ggtree")
# an R package for visualization of tree and annotation data
 library("ggtree")
if ( require("VGAM")         == FALSE ) install.packages("VGAM")
# Vector Generalized Linear and Additive Models
 library("VGAM")



# 1. Read and format data --------------------------------------------------

# Read shorbirds.csv file (Morphology & behaviour data)
Shorebird_data <- read.csv("shorebird_data.csv", header = TRUE, sep = ",",
stringsAsFactors=TRUE)
 # Use Species_ID as row labels
row.names(Shorebird_data) <- Shorebird_data[, "Species_ID"]
 # Compute tongue/beak length ratio (ln-transformed)
Shorebird_data$Tongue_beak_ln_ratio <- log(Shorebird_data$Tongue_length /
Shorebird_data$Beak_length)
 # Weight scales as a volume, so transform it with a cubic root
Shorebird_data$Cubicroot_weight <- Shorebird_data$Weight ^(1/3)

# Read shorebirds_tree.tre file (Phylogenetic tree)
Phylogenetic_tree <- ape::read.tree("shorebirds_tree.tre")

# Read species silhouette images to be added on graphics; they must be present
in the "Species_silhouettes" subdirectory of the working directory
 # those named "-cropped" have been cropped from originals to remove white
margins using this tool: https://svgcrop.com/
Numenius_arquata       <- svgparser::read_svg("Species_silhouettes/
Numenius_arquata-cropped.svg")       # original by Rebecca Groom based on Photo
by Andreas Trepte (https://www.phylopic.org/images/910114e1-39c6-4eae-87d5-
fd702dc47161/numenius-arquata)
Charadrius             <- svgparser::read_svg("Species_silhouettes/Charadrius-
cropped.svg")             # original by Ferran Sayol (https://www.phylopic.org/
images/2ff4c7f3-d403-407d-a430-e0e2bc54fab0/charadrius)
Haematopus_ostralegus  <- svgparser::read_svg("Species_silhouettes/
Haematopus_ostralegus-cropped.svg")  # original by Wouter Koch (https://
www.phylopic.org/images/e485635e-7dc3-4cb5-afcd-fe21c6767c71/haematopus-
ostralegus)
Recurvirostra_avosetta <- svgparser::read_svg("Species_silhouettes/
Recurvirostra_avosetta-cropped.svg") # original by Alexandre Vong (https://
www.phylopic.org/images/3266d069-f33e-4f80-9d0a-d8b0fddd1fe2/recurvirostra-
avosetta)
Vanellus_vanellus      <- svgparser::read_svg("Species_silhouettes/
Vanellus_vanellus-cropped.svg")      # original by Nina Skinner (https://
www.phylopic.org/images/4249a198-4b12-4878-a89b-699e47e95803/vanellus-vanellus)
Pluvialis_squatarola   <- svgparser::read_svg("Species_silhouettes/
Pluvialis_squatarola-cropped.svg")   # original by Emily Willoughby (https://
www.phylopic.org/images/a861cc3b-4940-4b80-84d7-5d1b07ba6a28/pluvialis-
squatarola)
Arenaria_interpres     <- svgparser::read_svg("Species_silhouettes/
Arenaria_interpres-cropped.svg")     # original by Rebecca Groom (https://
www.phylopic.org/images/6b77616d-7e99-488a-97e0-c6e919c62ec9/arenaria-interpres)
Limosa_limosa          <- svgparser::read_svg("Species_silhouettes/
Limosa_limosa-cropped.svg")          # original by Alexandre Vong (https://
www.phylopic.org/images/1fda4a62-111c-4e4c-9e12-ac0da0656cd9/limosa-limosa)
```

```
Gallinago_gallinago    <- svgparser::read_svg("Species_silhouettes/
Gallinago_gallinago-cropped.svg")    # original by Jamie Whitehouse (https://
www.phylopic.org/images/fba21da5-0dc4-44d0-a5ac-a9a2e024f46f/gallinago-
gallinago)
Calidris_pugnax        <- svgparser::read_svg("Species_silhouettes/
Calidris_pugnax-cropped.svg")        # original by Edwin Price (https://
www.phylopic.org/images/3813ebf5-b823-4f08-b05c-076fa09426e5/calidris-pugnax)
Actitis_hypoleucos     <- svgparser::read_svg("Species_silhouettes/
Actitis_hypoleucos-cropped.svg")     # original by Ferran Sayol (https://
www.phylopic.org/images/9e9780c0-baaf-4c83-a2d5-58c62a62135e/actitis-hypoleucos)
Limosa_lapponica       <- svgparser::read_svg("Species_silhouettes/
Limosa_lapponica-cropped.svg")       # original by Alexandre Vong (https://
www.phylopic.org/images/a6a38d55-71be-43f2-b781-9a845662de20/limosa-lapponica)
Tringa_flavipes        <- svgparser::read_svg("Species_silhouettes/
Tringa_flavipes-cropped.svg")        # original by Andy Wilson (https://
www.phylopic.org/images/1c9fb51b-5615-40fb-a2a8-f201fd61ac02/tringa-flavipes)
Himantopus_himantopus  <- svgparser::read_svg("Species_silhouettes/
Himantopus_himantopus.svg")          # original by Nicolas Schtickzelle for this
study
# Phylopic original image credits: https://www.phylopic.org/permalinks/
9e255cf6d53087e5625f92cbb5d6710e075a10881ae3417ddaea90fc27a7a28a



# 2. Phylogenetic analysis & create the body size indices (phylogenetically
controlled and raw) via Principal Component Analysis on 5 body metrics --------

# Perform phylogenetically controlled PCA -----------------------------------
Phylogenetic_PCA <- as.princomp( phytools::phyl.pca(Phylogenetic_tree,
Shorebird_data[,
c("Body_size","Wing_length","Tarsus_length","Toe_length","Cubicroot_weight")],
                                                     method="BM", mode="corr") )

# Extract loadings
Phylogenetic_loadings <- as.data.frame(Phylogenetic_PCA$loadings) %>%
                          rename(PC1=1, PC2=2, PC3=3, PC4=4, PC5=5) %>%
                           mutate(PC1 = -1 * PC1, PC2 = -1 * PC2)

# Extract species scores, change sign of PC1 & PC2, and use PC1 as
phylogenetically controlled global body size index;
Phylogenetic_scores <- as.data.frame(Phylogenetic_PCA$scores) %>%
                        rename(PC1=1, PC2=2, PC3=3, PC4=4, PC5=5) %>%
                         mutate(PC1 = -1 * PC1, PC2 = -1 * PC2)
Shorebird_data$Phylogenetic_body_size_index <- Phylogenetic_scores$PC1
Shorebird_data$Phylogenetic_PC2             <- Phylogenetic_scores$PC2

# Plot the phylogenetic tree
 # Add suborder in tree
TMP <- full_join(as_tibble(Phylogenetic_tree), Shorebird_data[, c("Species_ID",
"Suborder")], by = c("label" = "Species_ID"))
Phylogenetic_tree2 <- treeio::as.treedata(TMP)
rm(TMP)
Phylogenetic_tree_plot <- ggtree::revts(ggtree(Phylogenetic_tree2, layout =
"rectangular")) +
                          scale_color_manual(values=c("orange","dodgerblue2") )
+
                          geom_tiplab(aes(color = Suborder), show.legend=FALSE,
align=TRUE, size=4, hjust=-0.2) +
                          geom_cladelabel(node=31, label="Scolopaci",
color="dodgerblue2", angle=90, offset=6, hjust=0.5, vjust=1.5, align=TRUE) +
                          geom_cladelabel(node=51, label="Charadrii",
color="orange", angle=90, offset=6, hjust=0.5, vjust=1.5, align=TRUE) +
                          annotate("text", x=4.5, y=-3, label="Millions
years\nago (Ma)", vjust=0, hjust=0.5, size=4) + coord_cartesian(ylim = c(0,29),
clip = "off") +
```

```
                           theme_tree2(axis.text.x=element_text(color="black"),
plot.margin = unit(c(t=0.3, r=0, b=1.2, l=0),"cm") )
rm(Phylogenetic_tree2)

# Plot PC1 and PC2 as color matrix corresponding to the phylogenetic tree
 # Format scores for display using geom_tile
Phylogenetic_scores_for_matrix_plot <-
reshape2::melt(as.matrix(Phylogenetic_scores[, c("PC1","PC2")])) %>%
                                        mutate(Var1=gsub("_"," ",Var1),
Var2=gsub("\\.","",Var2)) %>%
                                         mutate(value=round(value, 3))
 # Get species order in phylogenetic tree
Species_tree_order <- as.data.frame(get_taxa_name(Phylogenetic_tree_plot)) %>%
                         rename(Var1=1)%>%
                           mutate(order=row_number())
 # Reorder scores as in phylogenetic tree
Phylogenetic_scores_for_matrix_plot <- Phylogenetic_scores_for_matrix_plot %>%
                                        left_join(Species_tree_order,
by=join_by(Var1)) %>%
                                         arrange(order, Var2)
Phylogenetic_score_matrix_plot <-
ggplot(data=Phylogenetic_scores_for_matrix_plot, aes(Var2, reorder(Var1,
desc(order)), fill = value)) +
                                  geom_tile(color="black") +
                                  scale_fill_gradient2(low="deeppink",
mid="white", high="aquamarine", limit=c(-32,32), midpoint=0, space="Lab",
name=NULL) +
                                  coord_fixed() +
                                  theme_classic(base_size=18) +
                                  theme(axis.text=element_text(color="black"),
                                        axis.line.x=element_blank(),
axis.text.x=element_text(angle=90, vjust=0.5), axis.title.x=element_blank(),
axis.ticks.x=element_blank(),
                                        axis.line.y=element_blank(),
axis.text.y=element_blank(),                   axis.title.y=element_blank(),
axis.ticks.y=element_blank(),
                                        plot.margin =
unit(c(0.4,0,0.7,0),"cm") )
rm(Phylogenetic_scores_for_matrix_plot)
# Join tree plot and score matrix plot
Phylogenetic_tree_plot_with_phylogenetic_score_matrix <-
ggpubr::ggarrange( Phylogenetic_tree_plot, Phylogenetic_score_matrix_plot,

ncol=2, nrow=1, widths=c(0.8,0.2) )

# Scree plot (percentage of explained variance)
Phylogenetic_scree_plot <- factoextra::fviz_eig(Phylogenetic_PCA,
choice="variance",
                                                main="", ylim=c(0, 100),
geom=c("bar","line"), ylab="",
                                                barcolor="palegoldenrod",
barfill="palegoldenrod",
                                                addlabels=TRUE, labelsize=5,
hjust=0,

ggtheme=theme_classic(base_size=18) ) +
                            labs(x="Principal component (PC)", y="Explained
variance (%)") +
                            theme(axis.text=element_text(color="black"))

# Correlation circle plot
 # Create file to print customized labels
Phylogenetic_loadings_for_corr_circle <- Phylogenetic_loadings[, c("PC1","PC2")]
%>%
                                          mutate(name=gsub("_","
",rownames(Phylogenetic_loadings)))
```

```
Phylogenetic_corr_circle_plot <-
ggplot(data=Phylogenetic_loadings_for_corr_circle, aes(x=PC1, y=PC2)) +
                                  geom_vline(xintercept=0, linetype="dashed") +
                                  geom_hline(yintercept=0, linetype="dashed") +
                                  annotate("segment", x = 0, y = 0, xend =
Phylogenetic_loadings$PC1, yend = Phylogenetic_loadings$PC2,
                                           arrow = arrow(type = "closed", length
= unit(0.04, "npc"))) +
                                  geom_circle(aes(x0 = 0, y0 = 0, r = 1)) +
coord_fixed() +

geom_text(data=Phylogenetic_loadings_for_corr_circle, aes(x=PC1, y=PC2,
label=name),
                                            size=5,
nudge_x=c(-0.9,-1.2,-0.5,-1.2,-0.7), nudge_y=c(-0.02,0,0,0,0)) +
                                  theme_classic(base_size=18) +
                                  xlab("PC1") + ylab("PC2") +
                                  theme(axis.text=element_text(color="black"),
                                        axis.text.x=element_text(vjust=0.5), ,
                                        plot.margin = unit(c(2,0,0,0),"cm") )

# Loadings plot
 # Format loadings matrix for display using geom_tile and change sign of PC1
Phylogenetic_loadings_for_plot <-
reshape2::melt(as.matrix(Phylogenetic_loadings[, c("PC1","PC2")])) %>%
                                    mutate(Var1=gsub("_"," ",Var1), Var2=gsub("\
\.","",Var2)) %>%
                                     mutate(value=round(value, 2)) %>%
                                      arrange(Var1)
Phylogenetic_loadings_plot <- ggplot(data=Phylogenetic_loadings_for_plot,
aes(Var2, Var1, fill = value)) +
                               geom_tile(color="black") +
                               scale_fill_gradient2(low="blue", high="red",
mid="white",midpoint=0, limit=c(-1,1), space="Lab", name=NULL) +
                               coord_fixed() +
                               geom_text(aes(Var2, Var1, label=value),
color="black", size=5) +
                               theme_classic(base_size=20) +
                               theme(axis.text=element_text(color="black"),
                                     axis.text.x=element_text(angle=90,
vjust=0.5), axis.title.x=element_blank(), axis.title.y=element_blank(),
axis.ticks.x=element_blank(), axis.ticks.y=element_blank(),
                                     plot.margin = unit(c(2,0,1,0),"cm") )
rm(Phylogenetic_loadings_for_plot)

# Score plot
Phylogenetic_score_plot <-
ggplot(data=Shorebird_data[,c("Species","Species_ID","Suborder","Phylogenetic_bo
dy_size_index","Phylogenetic_PC2")], aes(x=Phylogenetic_body_size_index,
y=Phylogenetic_PC2, label=Species_ID)) +
                            geom_vline(xintercept=0, linetype="dashed") +
                            geom_hline(yintercept=0, linetype="dashed") +
                            scale_color_manual(values=c("orange","dodgerblue2") )
+
                            geom_point(aes(color=Suborder), size=3) +
                            geom_text_repel(aes(color=Suborder, size=3),
max.overlaps=6, show.legend=FALSE) +
                            # add species silhouettes
                            annotation_custom(Charadrius            , xmin=-22  ,
ymin=  1  , ymax=  1   +1.5*1.00) + # dubi & alex; height is proportional to
species size
                            annotation_custom(Arenaria_interpres    , xmin=-12  ,
ymin=- 6  , ymax=- 6   +1.5*1.25) + # inte; 1.25
                            annotation_custom(Actitis_hypoleucos    , xmin=-24.5,
ymin=- 3  , ymax=- 3   +1.5*1.37) + # hypo; 1.37
```

```
                            geom_segment(aes(x = -16.8, y = 0, xend = -20, yend
= -2)) +
                           annotation_custom(Calidris_pugnax       , xmin=- 5.8,
ymin=  2  , ymax=  2   +1.5*1.72) + # pugn; 1.72
                           annotation_custom(Gallinago_gallinago   , xmin=- 9  ,
ymin=- 3.2, ymax=- 3.2 +1.5*1.79) + # gall; 1.79
                           annotation_custom(Tringa_flavipes       , xmin=-14  ,
ymin=  3.1, ymax=  3.1 +1.5*1.80) + # glar; 1.80
                           annotation_custom(Pluvialis_squatarola  , xmin=- 1  ,
ymin=- 4.5, ymax=- 4.5 +1.5*1.80) + # squa; 1.80
                           annotation_custom(Haematopus_ostralegus , xmin= 15  ,
ymin=- 7  , ymax=- 7   +1.5*2.7 ) + # ostr; 2.70
                           annotation_custom(Limosa_lapponica      , xmin=  6  ,
ymin=- 7  , ymax=- 7   +1.5*2.62) + # lapp; 2.62
                           annotation_custom(Vanellus_vanellus     , xmin=  3  ,
ymin=- 2  , ymax=- 2   +1.5*2.70) + # vane; 2.70
                           annotation_custom(Limosa_limosa         , xmin=  8  ,
ymin=  4  , ymax=  4   +1.5*2.90) + # limo; 2.90
                           annotation_custom(Recurvirostra_avosetta, xmin= 16  ,
ymin=  2  , ymax=  2   +1.5*3.60) + # avos; 3.60
                           annotation_custom(Numenius_arquata      , xmin= 20  ,
ymin=- 5.7, ymax=- 5.7 +1.5*5.50) + # arqu; 5.50
                           annotation_custom(Himantopus_himantopus , xmin= 12  ,
ymin= 10  , ymax= 10   +1.5*3.50) + # hima; 3.50
                           theme_classic(base_size=18) +
                           xlab("PC1 = Phylogenetically controlled global body
size index") + ylab("PC2") +
                           theme(axis.text=element_text(color="black"),
                                 legend.position = "inside",
legend.position.inside=c(0.1,0.9), legend.text=element_text(size=18),
legend.box.background=element_rect(color="black", size=2),
                                 plot.margin = unit(c(0.5,0,0,0),"cm") )


### Arrange plots in a composite figure (Fig 1) -------------------------
Left_column  <-
ggpubr::ggarrange( Phylogenetic_tree_plot_with_phylogenetic_score_matrix,
                                   Phylogenetic_score_plot,
                                   ncol=1, nrow=2, heights=c(1,1.5),
                                   labels=c("(A) Phylogenetic tree with species
scores", "(B) Phylomorphospace with species scores"),
                                   font.label=list(size=18, color="black",
face="bold", family=NULL), label.x=0, hjust=0, vjust=0.5 )
Right_column <- ggpubr::ggarrange( Phylogenetic_scree_plot,
                                   Phylogenetic_corr_circle_plot,
                                   Phylogenetic_loadings_plot,
                                   ncol=1, nrow=3, heights=c(1,1,1),
                                   labels=c("(C) Scree plot", "(D) Correlation
circle", "(E) Loadings"),
                                   font.label=list(size=18, color="black",
face="bold", family=NULL), label.x=0, hjust=0, vjust=c(0.85,3,4) )
Figure_1 <- ggpubr::ggarrange( Left_column, Right_column,
                               ncol=2, nrow=1, widths=c(0.7,0.3) ) +
            theme(plot.margin=margin(1,1,1,1,"cm"))
 # Opening the graphical device to save it as pdf
pdf("Figure_1.pdf", width=18, height=14)
Figure_1
 # Closing the graphical device
dev.off()



# 3. Explain morphology by body size index: computing relative-to-expected (RTE)
morphology traits --------

## 3.A. Eye diameter ----------------------------------------------------
```

```
# Fit linear regression model
Eye_diameter_regression <- lm(Eye_diameter ~ Phylogenetic_body_size_index,
data=Shorebird_data, na.action=na.exclude)

# Calculate relative-to-expected (RTE) value
Shorebird_data$Eye_diameter_expected <- predict(Eye_diameter_regression)
Shorebird_data$RTE_Eye_diameter <- log(Shorebird_data$Eye_diameter /
Shorebird_data$Eye_diameter_expected)

# Create plot of fitted model
Eye_diameter_regression_plot <- ggplot(data=subset(Shorebird_data, !
is.na(Eye_diameter)), aes(x=Phylogenetic_body_size_index, y=Eye_diameter) ) +

scale_color_manual(values=c("orange","dodgerblue2") ) +
                                geom_point(aes(color=Suborder), size=2.5,
show.legend=TRUE) +
                                geom_smooth(method="lm", formula="y~x",
se=FALSE, show.legend=FALSE, color="black") +
                                ggpubr::stat_cor(            label.x=-0.2,
label.y=8.22, size=5, show.legend=FALSE) +
                                ggpubr::stat_regline_equation(label.x=-0.2,
label.y=7.5, size=5, show.legend=FALSE) +
                                theme_classic(base_size=18) +
                                xlab("Global body size index") + ylab("Eye
Diameter (mm)") +
                                theme(axis.text=element_text(color="black"),
                                      legend.position = "inside",
legend.position.inside=c(0.3,0.85), legend.text=element_text(size=18),
legend.box.background=element_rect(color="black", size=2))

# Perform T-test on suborders
Eye_diameter_t_test <- Shorebird_data %>%
                         rstatix::t_test(RTE_Eye_diameter ~ Suborder,
var.equal=TRUE) %>%
                           rstatix::add_significance()

# Create violin plot per surborder
Eye_diameter_violin_plot <- ggplot(data=subset(Shorebird_data, !
is.na(RTE_Eye_diameter)), aes(x=RTE_Eye_diameter, y=Suborder, fill=Suborder),
show.legend=FALSE) +
                            geom_violin(trim=FALSE, show.legend=FALSE) +
                            scale_fill_manual(values=c("orange","dodgerblue2"))
+
                            stat_summary(fun=mean, geom="point", shape=3,
size=12, show.legend=FALSE) +
                            geom_point(size=2, show.legend=FALSE) +
                            geom_vline(xintercept=0, linetype="dashed") +
                            theme_classic(base_size=18)  +
                            xlab("RTE eye diameter") +
                            theme(axis.text=element_text(color="black"),
axis.text.x=element_text(vjust=0, hjust=0.5), axis.text.y=element_text(angle=90,
vjust=0, hjust=0.5)) +
                            annotate(geom="text", size=5, x=-0.61, y=1.45,
hjust=0, label=paste0("t-test = ", round(Eye_diameter_t_test$statistic,
digits=3))) +
                            annotate(geom="text", size=5, x=-0.61, y=1.25,
hjust=0, label=paste0("df = ", Eye_diameter_t_test$df, ", p = ",
round(Eye_diameter_t_test$p, digits=3)))


## 3.B. Number of sensory pits ---------------------------------------------

# Fit linear regression model
Sensory_pits_regression <- lm(Sensory_pits ~ Phylogenetic_body_size_index,
data=Shorebird_data, na.action=na.exclude)
```

```
# Calculate relative-to-expected (RTE) value
Shorebird_data$Sensory_pits_expected <- predict(Sensory_pits_regression)
Shorebird_data$RTE_Sensory_pits <-
suppressWarnings( (log(Shorebird_data$Sensory_pits /
Shorebird_data$Sensory_pits_expected)))  # A NaN value is produced for
Numenius_arquata because Sensory_pits_expected is negative

# Create plot of fitted model
Sensory_pits_regression_plot <- ggplot(data=subset(Shorebird_data, !
is.na(Sensory_pits)), aes(x=Phylogenetic_body_size_index, y=Sensory_pits) ) +

scale_color_manual(values=c("orange","dodgerblue2") ) +
                                geom_point(aes(color=Suborder), size=2.5,
show.legend=FALSE) +
                                geom_smooth(method="lm", formula="y~x",
se=FALSE, show.legend=FALSE, color="black") +
                                ggpubr::stat_cor(             label.x=-5,
label.y= 4 , size=5, show.legend=FALSE) +
                                ggpubr::stat_regline_equation(label.x=-5,
label.y=-11, size=5, show.legend=FALSE) +
                                theme_classic(base_size=18) +
                                xlab("Global body size index") + ylab("Number of
sensory pits") +
                                theme(axis.text=element_text(color="black"))

# Perform T-test on suborders
Sensory_pits_t_test <- Shorebird_data %>%
                         rstatix::t_test(RTE_Sensory_pits ~ Suborder,
var.equal=TRUE) %>%
                           rstatix::add_significance()

# Create violin plot per surborder
Sensory_pits_violin_plot <- ggplot(data=subset(Shorebird_data, (!
is.na(RTE_Sensory_pits)) & (!is.nan(RTE_Sensory_pits))), aes(x=RTE_Sensory_pits,
y=Suborder, fill=Suborder), show.legend=FALSE) +
                            geom_violin(trim=FALSE, show.legend=FALSE) +
                            scale_fill_manual(values=c("orange","dodgerblue2"))
+
                            stat_summary(fun=mean, geom="point", shape=3,
size=12, show.legend=FALSE) +
                            geom_point(size=2, show.legend=FALSE) +
                            geom_vline(xintercept=0, linetype="dashed") +
                            theme_classic(base_size=18) +
                            xlab("RTE number of sensory pits") + ylab("") +
                            theme(axis.text=element_text(color="black"),
axis.text.y=element_text(angle=90, vjust=0, hjust=0.5)) +
                            annotate(geom="text", size=5, x=-2, y=1.7, hjust=0,
label = paste0("t-test = ", round(Sensory_pits_t_test$statistic, digits=3))) +
                            annotate(geom="text", size=5, x=-2, y=1.5, hjust=0,
label = paste0("df = ", Sensory_pits_t_test$df, ", p = ",
round(Sensory_pits_t_test$p, digits=3)))


## 3.C. Beak length ---------------------------------------------------------

# Fit linear regression model
Beak_length_regression <- lm(Beak_length ~ Phylogenetic_body_size_index,
data=Shorebird_data, na.action=na.exclude)

# Calculate relative-to-expected (RTE) value
Shorebird_data$Beak_length_expected <- predict(Beak_length_regression)
Shorebird_data$RTE_Beak_length <- log(Shorebird_data$Beak_length /
Shorebird_data$Beak_length_expected)

# Create plot of fitted model
```

```
Beak_length_regression_plot <- ggplot(data=subset(Shorebird_data, !
is.na(Beak_length)), aes(x=Phylogenetic_body_size_index, y=Beak_length) ) +

scale_color_manual(values=c("orange","dodgerblue2") ) +
                                geom_point(aes(color=Suborder), size=2.5,
show.legend=FALSE) +
                                geom_smooth(method="lm", formula="y~x", se=FALSE,
show.legend=FALSE, color="black") +
                                ggpubr::stat_cor(             label.x=-0.1,
label.y=17, size=5, show.legend=FALSE) + # to get R² instead of R + p, add:
aes(label = after_stat(rr.label)),
                                ggpubr::stat_regline_equation(label.x=-0.1,
label.y=9, size=5, show.legend=FALSE) +
                                theme_classic(base_size=18) +
                                xlab("Global body size index") + ylab("Beak
length (mm)") +
                                theme(axis.text=element_text(color="black"))

# Perform T-test on suborders
Beak_length_t_test <- Shorebird_data %>%
                         rstatix::t_test(RTE_Beak_length ~ Suborder,
var.equal=TRUE) %>%
                           rstatix::add_significance()

# Create violin plot per surborder
Beak_length_violin_plot <- ggplot(data=subset(Shorebird_data, !
is.na(RTE_Beak_length)), aes(x=RTE_Beak_length, y=Suborder, fill=Suborder),
show.legend=FALSE) +
                            geom_violin(trim=FALSE, show.legend=FALSE) +
                            scale_fill_manual(values=c("orange","dodgerblue2")) +
                            stat_summary(fun=mean, geom="point", shape=3,
size=12, show.legend=FALSE) +
                            geom_point(size=2, show.legend=FALSE) +
                            geom_vline(xintercept=0, linetype="dashed") +
                            theme_classic(base_size=18) +
                            xlab("RTE beak length") + ylab("") +
                            theme(axis.text=element_text(color="black"),
axis.text.y=element_text(angle=90, vjust=0, hjust=0.5)) +
                            annotate(geom="text", size=5, x=-1.32, y=1.7,
hjust=0, label=paste0("t-test = ", round(Beak_length_t_test$statistic,
digits=3))) +
                            annotate(geom="text", size=5, x=-1.32, y=1.5,
hjust=0, label=paste0("df = ", Beak_length_t_test$df, ", p = ",
round(Beak_length_t_test$p, digits=3)))


## 3.D. Tongue/beak length ratio -------------------------------------------

# Fit linear regression model
Tongue_beak_ln_ratio_regression <- lm(Tongue_beak_ln_ratio ~
Phylogenetic_body_size_index, data=Shorebird_data, na.action=na.exclude)

# Calculate relative-to-expected (RTE) value
Shorebird_data$Tongue_beak_ln_ratio_expected <-
predict(Tongue_beak_ln_ratio_regression)
Shorebird_data$RTE_Tongue_beak_ln_ratio <-
log(Shorebird_data$Tongue_beak_ln_ratio /
Shorebird_data$Tongue_beak_ln_ratio_expected)

# Create plot of fitted model
Tongue_beak_ln_ratio_regression_plot <- ggplot(data=subset(Shorebird_data, !
is.na(Tongue_beak_ln_ratio)), aes(x=Phylogenetic_body_size_index,
y=Tongue_beak_ln_ratio,) ) +

scale_color_manual(values=c("orange","dodgerblue2") ) +
```

```
                                        geom_point(aes(color=Suborder),
size=2.5, show.legend=FALSE) +
                                        geom_smooth(method="lm", formula="y~x",
se=FALSE, show.legend=FALSE, color="black") +

ggpubr::stat_cor(            label.x=-5, label.y=-1.38, size=5,
show.legend=FALSE) +

ggpubr::stat_regline_equation(label.x=-5, label.y=-1.48 , size=5,
show.legend=FALSE) +
                                        theme_classic(base_size=18) +
                                        xlab("Global body size index") +
ylab("Tongue/beak length ln-ratio")

theme(axis.text=element_text(color="black"))

# Perform T-test on suborders
Tongue_beak_ln_ratio_t_test <- Shorebird_data %>%
                                 rstatix::t_test(RTE_Tongue_beak_ln_ratio ~
Suborder, var.equal=TRUE) %>%
                                   rstatix::add_significance()

# Create violin plot per surborder
Tongue_beak_ln_ratio_violin_plot <- ggplot(data=subset(Shorebird_data, !
is.na(RTE_Tongue_beak_ln_ratio)), aes(x=RTE_Tongue_beak_ln_ratio, y=Suborder,
fill=Suborder), show.legend=FALSE) +
                                    geom_violin(trim=FALSE, show.legend=FALSE) +

scale_fill_manual(values=c("orange","dodgerblue2")) +
                                    stat_summary(fun=mean, geom="point",
shape=3, size=12, show.legend=FALSE) +
                                    geom_point(size=2, show.legend=FALSE) +
                                    geom_vline(xintercept=0, linetype="dashed")
+
                                    theme_classic(base_size=18) +
                                    xlab("RTE tongue/beak length ln-ratio") +
ylab("") +
                                    theme(axis.text=element_text(color="black"),
axis.title.x=element_text(hjust=0.7), axis.text.y=element_text(angle=90,
vjust=0, hjust=0.5)) +
                                    annotate(geom="text", size=5, x=-2.25,
y=1.6, hjust=0, label=paste0("t-test = ",
round(Tongue_beak_ln_ratio_t_test$statistic, digits=3))) +
                                    annotate(geom="text", size=5, x=-2.25,
y=1.4, hjust=0, label=paste0("df = ", Tongue_beak_ln_ratio_t_test$df, ", p = ",
round(Tongue_beak_ln_ratio_t_test$p, digits=3)))

### Arrange plots in a composite figure (Fig 2) -------------------------
Figure_2 <- ggpubr::ggarrange(Eye_diameter_regression_plot,
Sensory_pits_regression_plot, Beak_length_regression_plot,
Tongue_beak_ln_ratio_regression_plot,
                              NULL,                        NULL,
NULL,                        NULL,
                              Eye_diameter_violin_plot,
Sensory_pits_violin_plot,     Beak_length_violin_plot,
Tongue_beak_ln_ratio_violin_plot,
                              ncol=4, nrow=3, heights=c(1,0.05,1), align="hv",
                              labels=c("(A) Eye (21 species)", "(B) Sensory pits
(7 species)", "(C) Beak (28 species)", "(D) Tongue (18 species)",
                                       "",       "",               "",
""),
                              font.label=list(size=18, color="black",
face="bold", family=NULL), hjust=c(0,0,0,0), vjust=c(-0.3,-0.3,-0.3,-0.3)) +
            theme(plot.margin=margin(2,1,1,1,"cm"))
 # Opening the graphical device to save it as pdf
pdf("Figure_2.pdf", width=18, height=10)
```

```
Figure_2
 # Closing the graphical device
dev.off()



# 4. Explore relations among morphological factors and with feeding behaviours
--------

## 4.A. Fit linear regression model between RTE nb of sensory pits & eye
diameter --------
RTENSP_RTEED_regression_plot <- ggplot(data=subset(Shorebird_data, !
is.na(RTE_Eye_diameter) & (!is.na(RTE_Sensory_pits))), aes(x=RTE_Eye_diameter,
y=RTE_Sensory_pits) ) +

scale_color_manual(values=c("orange","dodgerblue2") ) +
                                geom_smooth(method="lm", formula="y~x",
se=FALSE, show.legend=FALSE, color="black") +
                                geom_point(aes(color=Suborder), size=2.5,
show.legend=TRUE) +
                                ggpubr::stat_cor(             label.x=-0.25,
label.y=-1.38, size=5, show.legend=FALSE) +
                                ggpubr::stat_regline_equation(label.x=-0.25,
label.y=-1.55, size=5, show.legend=FALSE) +
                                theme_classic(base_size=18) +
                                xlab("RTE eye diameter") + ylab("RTE number of
sensory pits") +
                                theme(axis.text=element_text(color="black"),
                                      legend.position = "inside",
legend.position.inside=c(0.8,0.9), legend.text=element_text(size=18),
legend.box.background=element_rect(color="black", size=2),
                                      plot.margin = unit(c(1,1,0,0),"cm") )


### Arrange plot in a figure (Fig 3) -------------------------
Figure_3 <- ggpubr::ggarrange(RTENSP_RTEED_regression_plot,
                              ncol=1, nrow=1,
                              font.label=list(size=18, color="black",
face="bold", family=NULL)) +
                              theme(plot.margin=margin(1,1,1,1,"cm"))
 # Opening the graphical device to save it as pdf
pdf("Figure_3.pdf", width=5, height=5)
Figure_3
 # Closing the graphical device
dev.off()


## 4.B. Fit binomial logistic regression model between Locomotion behaviour &
RTE beak length --------

# Create 0,1 dummy var for Locomotion (needed for geom-smooth)
Shorebird_data <- Shorebird_data %>%
                    mutate(Locomotion_numeric=case_when(Locomotion=="CW"~1,
Locomotion=="SRS"~0) )

# Fit logistic regression model
Locomotion_RTEBL_logistic_regression <- glm(Locomotion_numeric ~
RTE_Beak_length, data=Shorebird_data, family=binomial(link="logit"))
Locomotion_RTEBL_logistic_regression_results <-
summary(Locomotion_RTEBL_logistic_regression)
Locomotion_RTEBL_logistic_regression_slope <-
as.data.frame(Locomotion_RTEBL_logistic_regression_results$coefficients)
Locomotion_RTEBL_logistic_regression_slope <-
Locomotion_RTEBL_logistic_regression_slope %>%
```

```
filter(row.names(Locomotion_RTEBL_logistic_regression_slope) ==
"RTE_Beak_length")

# Create plot of fitted model
Locomotion_RTEBL_regression_plot <- ggplot(data=Shorebird_data,
aes(x=RTE_Beak_length, y=Locomotion_numeric) ) + # To fit a logistic regression,
you need to coerce the values to a numeric vector lying between 0 and 1

scale_color_manual(values=c("orange","dodgerblue2")) +
                                    geom_smooth(method="glm", formula="y~x",
method.args=list(family="binomial"), se=FALSE, show.legend=FALSE, color="black")
+
                                    geom_point(aes(color=Suborder), size=2.5,
stroke = 2, shape = 21, show.legend=TRUE) +
                                    annotate(geom="text", size=5, x=-1, y=0.9,
hjust=0, label=paste0("z = ",
round(Locomotion_RTEBL_logistic_regression_slope$"z value", digits=3),", p = ",
round(Locomotion_RTEBL_logistic_regression_slope$"Pr(>|z|)",digits=3))) +
                                    theme_classic(base_size=18) +
                                    xlab("RTE beak length") + ylab("Locomotion
behaviour") +
                                    scale_y_continuous(breaks = c(0,1)) +
                                    theme(axis.text.y=element_blank()) +
                                    annotate(geom="text", size=5, x=-1, y=0,
hjust=1.7, label="SRS") + annotate(geom="text", size=5, x=-1, y=1, hjust=1.85,
label="CW") +
                                    coord_cartesian(xlim=c(-1,0.5), clip='off')
+
                                    theme(axis.text=element_text(color="black"),
                                          legend.position = "inside",
legend.position.inside=c(0.8,0.5), legend.text=element_text(size=18),
legend.box.background=element_rect(color="black", size=2),
                                          plot.margin = unit(c(1,1,0,0),"cm") )


## 4.C. Fit cumulative logistic regression model between Capture behaviour & RTE
beak length --------
 # NB: Order of ordinal capture behaviours: PE < PE-PR < PR

# Fit logistic regression model
Capture_RTEBL_cumulative_logistic_regression <-
suppressWarnings( VGAM::vglm(as.ordered(Capture) ~ RTE_Beak_length, # warnings
are concerned with deprecated use of "logit" instead of logitlink

data = Shorebird_data,

family = cumulative(link = "logit", parallel = TRUE, reverse = TRUE)) )
Capture_RTEBL_Estimates <-
suppressWarnings( as.data.frame(coef(summary(Capture_RTEBL_cumulative_logistic_r
egression))) )
Capture_RTEBL_Slope <- Capture_RTEBL_Estimates["RTE_Beak_length",]

# Compute predicted probabilities for observed species + regularly spaced x
values for plotting fitted curves
Regular_points_RTEBL <-
data.frame("RTE_Beak_length"=seq(min(Shorebird_data[,"RTE_Beak_length"]),max(Sho
rebird_data[,"RTE_Beak_length"]), by = 0.01))
Capture_RTEBL_proba <- suppressWarnings(

as.data.frame(predictvglm(Capture_RTEBL_cumulative_logistic_regression, type =
"response")) %>% # observed values
                              bind_cols(Shorebird_data[, c("RTE_Beak_length",
"Suborder", "Capture")])  %>%
```

```
                              bind_rows(as.data.frame(cbind(Regular_points_RTEBL,
predictvglm(Capture_RTEBL_cumulative_logistic_regression, Regular_points_RTEBL,
type = "response")))) %>% # regularly spaced x values for plotting fitted curves
                                 rename("Proba_Capture_PE"="PE",
"Proba_Capture_PE_PR"="PE-PR", "Proba_Capture_PR"="PR") %>%
                                    group_by(Capture) %>%
                                      mutate(Proba_Capture_PE
=case_when(Capture!="PE"     ~ NA, TRUE ~ Proba_Capture_PE   ),

Proba_Capture_PE_PR=case_when(Capture!="PE-PR" ~ NA, TRUE ~
Proba_Capture_PE_PR),
                                             Proba_Capture_PR
=case_when(Capture!="PR"     ~ NA, TRUE ~ Proba_Capture_PR   ))  )

# Create plot of fitted model
Capture_RTEBL_regression_plot <- ggplot(data=Capture_RTEBL_proba) +

geom_line(data=Capture_RTEBL_proba[is.na(Capture_RTEBL_proba$Suborder),],
aes(x=RTE_Beak_length, y=Proba_Capture_PE   ), color="aquamarine2", linewidth=1,
show.legend=FALSE, na.rm=TRUE ) +

geom_line(data=Capture_RTEBL_proba[is.na(Capture_RTEBL_proba$Suborder),],
aes(x=RTE_Beak_length, y=Proba_Capture_PE_PR), color="deeppink",   linewidth=1,
show.legend=FALSE, na.rm=TRUE ) +

geom_line(data=Capture_RTEBL_proba[is.na(Capture_RTEBL_proba$Suborder),],
aes(x=RTE_Beak_length, y=Proba_Capture_PR   ), color="purple",  linewidth=1,
show.legend=FALSE, na.rm=TRUE ) +

scale_color_manual(values=c("orange","dodgerblue2","white")) +
                                  geom_point(data=Capture_RTEBL_proba[!
is.na(Capture_RTEBL_proba$Suborder),], aes(x=RTE_Beak_length,
y=Proba_Capture_PE,    color=Suborder), size=2.5, stroke = 2, shape = 21,
show.legend=FALSE, na.rm=TRUE ) +
                                  geom_point(data=Capture_RTEBL_proba[!
is.na(Capture_RTEBL_proba$Suborder),], aes(x=RTE_Beak_length,
y=Proba_Capture_PE_PR, color=Suborder), size=2.5, stroke = 2, shape = 21,
show.legend=FALSE, na.rm=TRUE ) +
                                  geom_point(data=Capture_RTEBL_proba[!
is.na(Capture_RTEBL_proba$Suborder),], aes(x=RTE_Beak_length,
y=Proba_Capture_PR,    color=Suborder), size=2.5, stroke = 2, shape = 21,
show.legend=FALSE, na.rm=TRUE ) +
                                  annotate(geom="text", size=6, x=-0.85, y=0.8 ,
hjust=0, label="Pecking", color="aquamarine2") +
                                  annotate(geom="text", size=6, x=-0.85, y=0.1 ,
hjust=0, label="Both"   , color="deeppink") +
                                  annotate(geom="text", size=6, x= 0.2 , y=0.85,
hjust=0, label="Probing", color="purple") +
#                                  annotate(geom="text", size=5, x=-0.95, y=0.6 ,
hjust=0, label=paste0("Slope = ", round(Capture_RTEBL_Slope$Estimate,
digits=3))) +
                                  annotate(geom="text", size=5, x=-0.95, y=0.5 ,
hjust=0, label=paste0("z = ",      round(Capture_RTEBL_Slope$"z value",
digits=3))) +
                                  annotate(geom="text", size=5, x=-0.95, y=0.4 ,
hjust=0, label=paste0("p = ",      round(Capture_RTEBL_Slope$"Pr(>|z|)",
digits=3))) +
                                  theme_classic(base_size=18) +
                                  xlab("RTE beak length") + ylab("Capture
behaviour probability") +
                                  ylim(0,1) +
                                  theme(axis.text=element_text(color="black"),
plot.margin = unit(c(1,1,0,0),"cm") )
```

```
## 4.D. Fit multinomial logistic regression model between Transport behaviour & RTE beak length --------

# Fit logistic regression model & null model
Transport_RTEBL_multinomial_logistic_regression <-  VGAM::vglm(Transport ~ RTE_Beak_length,
                                                                  data = Shorebird_data,
                                                                  family = multinomial )
Transport_RTEBL_multinomial_logistic_regression_null_model <- VGAM::vglm(Transport ~  1,
                                                                  data = Shorebird_data,
                                                                  family = multinomial )
Transport_RTEBL_multinomial_logistic_regression_Estimates <- as.data.frame(coef(summary(Transport_RTEBL_multinomial_logistic_regression)))

# likelihood ration test of the global effect of RTE_Beak_length
Transport_RTEBL_multinomial_logistic_regression_DF_model      <- Transport_RTEBL_multinomial_logistic_regression@df.residual
Transport_RTEBL_multinomial_logistic_regression_DF_null_model <- Transport_RTEBL_multinomial_logistic_regression_null_model@df.residual
Transport_RTEBL_multinomial_logistic_regression_LL_model      <- Transport_RTEBL_multinomial_logistic_regression@criterion$loglikelihood
Transport_RTEBL_multinomial_logistic_regression_LL_null_model <- Transport_RTEBL_multinomial_logistic_regression_null_model@criterion$loglikelihood
Transport_RTEBL_multinomial_logistic_regression_LL_diff       <- Transport_RTEBL_multinomial_logistic_regression_LL_model      - Transport_RTEBL_multinomial_logistic_regression_LL_null_model
Transport_RTEBL_multinomial_logistic_regression_DF_diff       <- Transport_RTEBL_multinomial_logistic_regression_DF_null_model - Transport_RTEBL_multinomial_logistic_regression_DF_model
Transport_RTEBL_multinomial_logistic_regression_p             <- 1 - pchisq(Transport_RTEBL_multinomial_logistic_regression_LL_diff, df=Transport_RTEBL_multinomial_logistic_regression_DF_diff)

# Compute predicted probabilities for observed species + regularly spaced x values for plotting fitted curves
Transport_RTEBL_proba <- as.data.frame(predictvglm(Transport_RTEBL_multinomial_logistic_regression, type = "response")) %>% # observed values
                           bind_cols(Shorebird_data[, c("RTE_Beak_length", "Suborder", "Transport")])  %>%
                             bind_rows(as.data.frame(cbind(Regular_points_RTEBL, predictvglm(Transport_RTEBL_multinomial_logistic_regression, Regular_points_RTEBL, type = "response")))) %>% # regularly spaced x values for plotting fitted curves
                               rename("Proba_Transport_BA"="BA", "Proba_Transport_BA_ST"="BA-ST", "Proba_Transport_LI"="LI") %>%
                                 group_by(Transport) %>%
                                   mutate(Proba_Transport_BA    =case_when(Transport!="BA"    ~ NA, TRUE ~ Proba_Transport_BA   ),
                                          Proba_Transport_BA_ST=case_when(Transport!="BA-ST" ~ NA, TRUE ~ Proba_Transport_BA_ST),
                                          Proba_Transport_LI    =case_when(Transport!="LI"    ~ NA, TRUE ~ Proba_Transport_LI   )  )

# Create plot of fitted model
Transport_RTEBL_regression_plot <- ggplot(data=Transport_RTEBL_proba) +
  geom_line(data=Transport_RTEBL_proba[is.na(Transport_RTEBL_proba$Suborder),],
```

```
aes(x=RTE_Beak_length, y=Proba_Transport_BA   ), color="aquamarine2",
linewidth=1, show.legend=FALSE, na.rm=TRUE ) +

geom_line(data=Transport_RTEBL_proba[is.na(Transport_RTEBL_proba$Suborder),],
aes(x=RTE_Beak_length, y=Proba_Transport_BA_ST), color="deeppink",
linewidth=1, show.legend=FALSE, na.rm=TRUE ) +

geom_line(data=Transport_RTEBL_proba[is.na(Transport_RTEBL_proba$Suborder),],
aes(x=RTE_Beak_length, y=Proba_Transport_LI   ), color="purple",  linewidth=1,
show.legend=FALSE, na.rm=TRUE ) +

scale_color_manual(values=c("orange","dodgerblue2","white")) +
                                    geom_point(data=Transport_RTEBL_proba[!
is.na(Transport_RTEBL_proba$Suborder),], aes(x=RTE_Beak_length,
y=Proba_Transport_BA,    color=Suborder), size=2.5, stroke = 2, shape = 21,
show.legend=FALSE, na.rm=TRUE ) +
                                    geom_point(data=Transport_RTEBL_proba[!
is.na(Transport_RTEBL_proba$Suborder),], aes(x=RTE_Beak_length,
y=Proba_Transport_BA_ST, color=Suborder), size=2.5, stroke = 2, shape = 21,
show.legend=FALSE, na.rm=TRUE ) +
                                    geom_point(data=Transport_RTEBL_proba[!
is.na(Transport_RTEBL_proba$Suborder),], aes(x=RTE_Beak_length,
y=Proba_Transport_LI,    color=Suborder), size=2.5, stroke = 2, shape = 21,
show.legend=FALSE, na.rm=TRUE ) +
                                    annotate(geom="text", size=6, x= 0.1, y=0.25,
hjust=0, label="Ballistic", color="aquamarine2") +
                                    annotate(geom="text", size=6, x= 0.1, y=0.55,
hjust=0, label="Ballistic\n+ surface\n   tension"   , color="deeppink") +
                                    annotate(geom="text", size=6, x=-0.6, y=1   ,
hjust=0, label="Lingual", color="purple") +
                                    annotate(geom="text", size=5, x=-0.95, y=0.5,
hjust=0, label=paste0("Chi² = ",
round(Transport_RTEBL_multinomial_logistic_regression_LL_diff, digits=3))) +
                                    annotate(geom="text", size=5, x=-0.95, y=0.4,
hjust=0, label=paste0("df = ",
Transport_RTEBL_multinomial_logistic_regression_DF_diff  )) +
                                    annotate(geom="text", size=5, x=-0.95, y=0.3,
hjust=0, label=paste0("p = ",
round(Transport_RTEBL_multinomial_logistic_regression_p,      digits=3))) +
                                    theme_classic(base_size=18) +
                                    xlab("RTE beak length") + ylab("Transport
behaviour probability") +
                                    ylim(0,1) +
                                    theme(axis.text=element_text(color="black"),
plot.margin = unit(c(1,1,0,0),"cm") )
rm (Regular_points_RTEBL)


## 4.E. Fit multinomial logistic regression model between Transport behaviour &
RTE_Tongue_beak_ln_ratio --------

# Fit logistic regression model & null model
Transport_RTETB_multinomial_logistic_regression <-  VGAM::vglm(Transport ~
RTE_Tongue_beak_ln_ratio,
                                                                 data =
Shorebird_data[!is.na(Shorebird_data$RTE_Tongue_beak_ln_ratio),],
                                                                 family =
multinomial )
Transport_RTETB_multinomial_logistic_regression_null_model <-
VGAM::vglm(Transport ~  1,
                                                                            data =
Shorebird_data[!is.na(Shorebird_data$RTE_Tongue_beak_ln_ratio),],
                                                                            family
= multinomial )
Transport_RTETB_multinomial_logistic_regression_Estimates <-
as.data.frame(coef(summary(Transport_RTETB_multinomial_logistic_regression)))
```

```
# likelihood ratio test of the global effect of RTE_Tongue_beak_ln_ratio
Transport_RTETB_multinomial_logistic_regression_DF_model      <-
Transport_RTETB_multinomial_logistic_regression@df.residual
Transport_RTETB_multinomial_logistic_regression_DF_null_model <-
Transport_RTETB_multinomial_logistic_regression_null_model@df.residual
Transport_RTETB_multinomial_logistic_regression_LL_model      <-
Transport_RTETB_multinomial_logistic_regression@criterion$loglikelihood
Transport_RTETB_multinomial_logistic_regression_LL_null_model <-
Transport_RTETB_multinomial_logistic_regression_null_model@criterion$loglikeliho
od
Transport_RTETB_multinomial_logistic_regression_LL_diff       <-
Transport_RTETB_multinomial_logistic_regression_LL_model      -
Transport_RTETB_multinomial_logistic_regression_LL_null_model
Transport_RTETB_multinomial_logistic_regression_DF_diff       <-
Transport_RTETB_multinomial_logistic_regression_DF_null_model -
Transport_RTETB_multinomial_logistic_regression_DF_model
Transport_RTETB_multinomial_logistic_regression_p             <- 1 -
pchisq(Transport_RTETB_multinomial_logistic_regression_LL_diff,
df=Transport_RTETB_multinomial_logistic_regression_DF_diff)

# Compute predicted probabilities for observed species + regularly spaced x
values for plotting fitted curves
Regular_points_RTETB <-
data.frame("RTE_Tongue_beak_ln_ratio"=seq(min(Shorebird_data[,
"RTE_Tongue_beak_ln_ratio"], na.rm=TRUE),max(Shorebird_data[,
"RTE_Tongue_beak_ln_ratio"], na.rm=TRUE), by = 0.01))
Transport_RTETB_proba <-
as.data.frame(predictvglm(Transport_RTETB_multinomial_logistic_regression, type
= "response")) %>% # observed values
                                          bind_cols(Shorebird_data[!
is.na(Shorebird_data$RTE_Tongue_beak_ln_ratio), c("RTE_Tongue_beak_ln_ratio",
"Suborder", "Transport")])  %>%

bind_rows(as.data.frame(cbind(Regular_points_RTETB,
predictvglm(Transport_RTETB_multinomial_logistic_regression,
Regular_points_RTETB, type = "response")))) %>% # regularly spaced x values for
plotting fitted curves
                                             rename("Proba_Transport_BA"="BA",
"Proba_Transport_BA_ST"="BA-ST", "Proba_Transport_LI"="LI") %>%
                                               group_by(Transport) %>%
                                                 mutate(Proba_Transport_BA
=case_when(Transport!="BA"    ~ NA, TRUE ~ Proba_Transport_BA   ),

Proba_Transport_BA_ST=case_when(Transport!="BA-ST" ~ NA, TRUE ~
Proba_Transport_BA_ST),
                                                       Proba_Transport_LI
=case_when(Transport!="LI"    ~ NA, TRUE ~ Proba_Transport_LI   )  )

# Create plot of fitted model
Transport_RTETB_regression_plot <- ggplot(data=Transport_RTETB_proba) +

geom_line(data=Transport_RTETB_proba[is.na(Transport_RTETB_proba$Suborder),],
aes(x=RTE_Tongue_beak_ln_ratio, y=Proba_Transport_BA   ), color="aquamarine2",
linewidth=1, show.legend=FALSE, na.rm=TRUE ) +

geom_line(data=Transport_RTETB_proba[is.na(Transport_RTETB_proba$Suborder),],
aes(x=RTE_Tongue_beak_ln_ratio, y=Proba_Transport_BA_ST), color="deeppink",
linewidth=1, show.legend=FALSE, na.rm=TRUE ) +

geom_line(data=Transport_RTETB_proba[is.na(Transport_RTETB_proba$Suborder),],
aes(x=RTE_Tongue_beak_ln_ratio, y=Proba_Transport_LI   ), color="purple",
linewidth=1, show.legend=FALSE, na.rm=TRUE ) +

scale_color_manual(values=c("orange","dodgerblue2","white")) +
```

```
                                    geom_point(data=Transport_RTETB_proba[!
is.na(Transport_RTETB_proba$Suborder),], aes(x=RTE_Tongue_beak_ln_ratio,
y=Proba_Transport_BA,    color=Suborder), size=2.5, stroke = 2, shape = 21,
show.legend=FALSE, na.rm=TRUE ) +
                                    geom_point(data=Transport_RTETB_proba[!
is.na(Transport_RTETB_proba$Suborder),], aes(x=RTE_Tongue_beak_ln_ratio,
y=Proba_Transport_BA_ST, color=Suborder), size=2.5, stroke = 2, shape = 21,
show.legend=FALSE, na.rm=TRUE ) +
                                    geom_point(data=Transport_RTETB_proba[!
is.na(Transport_RTETB_proba$Suborder),], aes(x=RTE_Tongue_beak_ln_ratio,
y=Proba_Transport_LI,    color=Suborder), size=2.5, stroke = 2, shape = 21,
show.legend=FALSE, na.rm=TRUE ) +
                                    annotate(geom="text", size=6, x=-1  , y=0.04
, hjust=0, label="Ballistic", color="aquamarine2") +
                                    annotate(geom="text", size=6, x=-1.1, y=0.85,
hjust=0, label="Ballistic + surface tension"   , color="deeppink") +
                                    annotate(geom="text", size=6, x=-1.2, y=0.25,
hjust=0, label="Lingual", color="purple") +
                                    annotate(geom="text", size=5, x= 0.4, y=0.5,
hjust=0, label=paste0("Chi² = ",
round(Transport_RTETB_multinomial_logistic_regression_LL_diff, digits=3))) +
                                    annotate(geom="text", size=5, x= 0.4, y=0.4,
hjust=0, label=paste0("df = ",
Transport_RTETB_multinomial_logistic_regression_DF_diff) ) +
                                    annotate(geom="text", size=5, x= 0.4, y=0.3,
hjust=0, label=paste0("p = ",
round(Transport_RTETB_multinomial_logistic_regression_p,       digits=3))) +
                                    theme_classic(base_size=18) +
                                    xlab("RTE tongue/beak length ln-ratio") +
ylab("Transport behaviour probability") +
                                    ylim(0,1) +
                                    theme(axis.text=element_text(color="black"),
plot.margin = unit(c(1,1,0,0),"cm") )
rm(Regular_points_RTETB)


### Arrange plots in a composite figure (Fig 4) ------------------------
Figure_4 <- ggpubr::ggarrange(Locomotion_RTEBL_regression_plot,
Capture_RTEBL_regression_plot, Transport_RTEBL_regression_plot,
Transport_RTETB_regression_plot,
                               ncol=4, nrow=1, align="hv",
                               labels=c("(A) Locomotion behaviour\n        ~ Beak
length",
                                        "(B) Capture behaviour\n        ~ Beak
length",
                                        "(C) Transport behaviour\n        ~ Beak
length",
                                        "(D) Transport behaviour\n        ~
Tongue/beak length"),
                               font.label=list(size=18, color="black",
face="bold", family=NULL), hjust=0, vjust=c(0.3,0.3,0.3,0.3)) +
            theme(plot.margin=margin(2,1,1,1,"cm"))
# Opening the graphical device to save it as pdf
pdf("Figure_4.pdf", width=18, height=5.5)
Figure_4
# Closing the graphical device
dev.off()
```

**Data S1. (separate file)**

*shorebird_data.csv*: spreadsheet with morphological and behavioral traits of the 29 shorebirds species:

- Species: species latin name
- Species_ID: species code used to label species on figures, made from the first 4 letters of the specific epithet
- Suborder: the suborder (Charadrii or Scolopaci) to which the species belongs
- Body_size: length (in cm) of the body, measured from tip of beak to tip of tail
- Wing_length: length (in mm) of the wing, measured from carpal joint to the tip of the longest primary
- Beak_length: length (in mm) of the beak, the chord of the culmen, measured from imputation of the feathers to the tip of the upper mandible
- Tarsus_length length (in mm) of the tarsus, measured from from the middle point of the joint between tibia and tarsus to the middle point of the joint between the tarsus and middle toe in front of the leg
- Toe_length length (in mm) of the toe, measured from this point to the tip of the middle claw
- Weight: body weight (in grams)
- Tongue_length: length (in mm) of the tongue, measured from the level of the tips of the posters-lateral papillae
- Eye_diameter: diameter (in mm) of the eye
- Sensory_pits: number of sensory pits on the beak
- Capture: food capture behavior according to Baguette et al. 2024: pecking (PE), probing (PR), pecking-probing (PE-PR),
- Locomotion: foraging locomotion behavior according to Baguette et al. 2024: stop-run-stop (SRS), continuous walking (CW)
- Transport: food capture behavior according to Baguette et al. 2024: lingual (LI), ballistic (BA) and ballistic-surface tension (BA-ST)

**Data S2. (separate file)**

*shorebirds_tree.tre*: phylogenetic tree information for the 29 shorebirds species, formatted as a text file to be read by the ape R package (*1*)
(using command "`ape::read.tree("shorebirds_tree.tre")`"
